# Biophysical mapping of TREM2-ligand interactions reveals shared surfaces for engagement of multiple AD ligands

**DOI:** 10.1101/2024.05.23.595592

**Authors:** Jessica A. Greven, Joshua R. Wydra, Rory A. Greer, Christopher Camitta, Yuhua Song, Tom J. Brett, Jennifer M. Alexander-Brett

## Abstract

TREM2 is a signaling receptor expressed on microglia that has emerged as an important potential drug target for Alzheimer’s disease and other neurodegenerative diseases. While a number of TREM2 signaling ligands have been identified, little is known regarding the structural details of how it engages them. To better understand this, we created a protein library of 28 different TREM2 variants and 11 different sTREM2 variants that could be used to map interactions with various ligands using biolayer interferometry (BLI). The variants are located in previously identified putative binding surfaces on TREM2 called the hydrophobic site, basic site, and site 2. We found that mutations to the hydrophobic site ablated binding to apoE4, oAb42, and TDP-43. Competition binding experiments further supported that apoE4 and oAb42 share overlapping binding sites on TREM2. In contrast, binding to IL-34 was mediated by the basic site at a surface centering on R76. Competition binding experiments validated a unique site for IL-34, showing little to no competition with either oAb42 or apoE4. Altogether, our results suggest that TREM2 utilizes the hydrophobic site (consisting of CDR1, CDR2, and CDR3) as a common site to engage multiple ligands, and further implies that pharmaceutical strategies targeting this surface might be effective to modulate TREM2 functions.

## Introduction

The triggering receptor expressed on myeloid cells-2 (TREM2) plays a central role in regulating myeloid cell maturation and function in the setting of numerous human pathologies, including neurodegenerative diseases, metabolic diseases, and cancers [1]. The potential importance of TREM2 in neuronal health was first indicated by large-scale genetic studies that identified rare TREM2 point variants, namely R47H and R62H, as risk factors for developing late-onset Alzheimer’s disease (AD) [2–4]. TREM2 is mainly expressed on macrophages and microglia, and responds to ligands associated with tissue damage. Within the setting of neurodegenerative diseases, particularly AD, responses signaled through TREM2 trigger activation of microglia into a Damage Associated Microglia (DAM) phenotype. This signaling can enhance protective functions of microglia, including chemotaxis, phagocytosis, suppression of inflammatory signaling, and boosting microglia survival and proliferation [5]. In addition, TREM2 can be proteolytically released from the cell surface or alternatively transcribed to produce soluble TREM2 (sTREM2), which also has beneficial functions in the setting of Alzheimer’s disease. For example, sTREM2 has been shown to reduce amyloid beta pathologies in AD mouse models [6], and increased sTREM2 levels in human CSF appear to correlate increased cognitive reserve [7]. In addition, sTREM2 has been shown to bind amyloid beta (Aβ) and inhibit its aggregation [8, 9], and can also disaggregate Aβ oligomers and filaments [10]. For these and other reasons, TREM2 has emerged as a viable drug target for Alzheimer’s disease, especially in the preclinical stage.

In recent years, a number of potential TREM2 ligands with relevance to AD have been identified. The first was apolipoprotein E (apoE) [11, 12], which also has an allelic variant, apoE4, that is a strong genetic risk factor for AD. ApoE has been shown to be a signaling ligand for TREM2 [13, 14] and its association with amyloid beta might assist the phagocytosis and degradation of amyloid beta oligomers by microglia via TREM2 [14]. Oligomeric amyloid beta (oAβ) is also a signaling ligand for TREM2 [6, 15, 16], with its engagement also triggering phagocytosis and degradation [15–18]. Quite recently, oligomeric TDP-43 was also identified as a phagocytosis-triggering ligand for TREM2 [19]. TDP-43 is the main component of insoluble aggregates found in patients with ALS [20], and these aggregates have also been identified in FTD and AD [21]. Additionally, the signaling cytokine IL-34 was recently identified as a TREM2 signaling ligand [22]. IL-34 has been linked to AD [23] and IL-34 mediated signaling in a microglial cell line was demonstrated [22]. Even though these potential ligands have been identified, the identity of which are the endogenous ligands of TREM2 in the setting of developing or established AD pathology is unknown.

Although a number of functional ligands for TREM2 have been identified, little is known regarding the structural mechanisms of their engagement. Such knowledge is critical not only to understand the structural basis for TREM2 signaling functions, but is also crucial to design therapeutic strategies that target TREM2. TREM2 is a single-pass transmembrane receptor consisting of an extracellular V-type Ig domain and intrinsically disordered stalk, transmembrane helix, and short cytoplasmic tail (**Fig. 1A**). This tail lacks any signal transduction or trafficking motifs, and TREM2 associates with the adaptor proteins DAP12 and DAP10 via transmembrane contacts to facilitate its trafficking and signaling. The Ig domain is likely the main region of TREM2 that engages extracellular ligands. We determined the first crystal structure of TREM2 and identified two potential ligand-binding surfaces on the Ig domain: a large hydrophobic site located on the distal end of the molecule in the CDR loops; and a large electropositive surface called the basic site that stretched partly around the midsection of the protein ([24] and **Fig. 1B**). Since then, there have been few studies showing how these surfaces might be involved in binding ligands. A co-crystal structure of TREM2 in complex with a soluble analog of the phospholipid phosphotidylserine showed that the phophatidylserine headgroup bound near the top of the basic site [25]. More recently, we showed that point mutations to the hydrophobic site diminished binding affinity for apoE4 almost 200-fold, suggesting that this protein mainly utilized the hydrophobic site on TREM2 for engagement [9]. In addition, a recent study used crosslinking and mass spectrometry to identify another region on TREM2 called site 2 that might engage oAβ42 [8].

**Fig 1:**
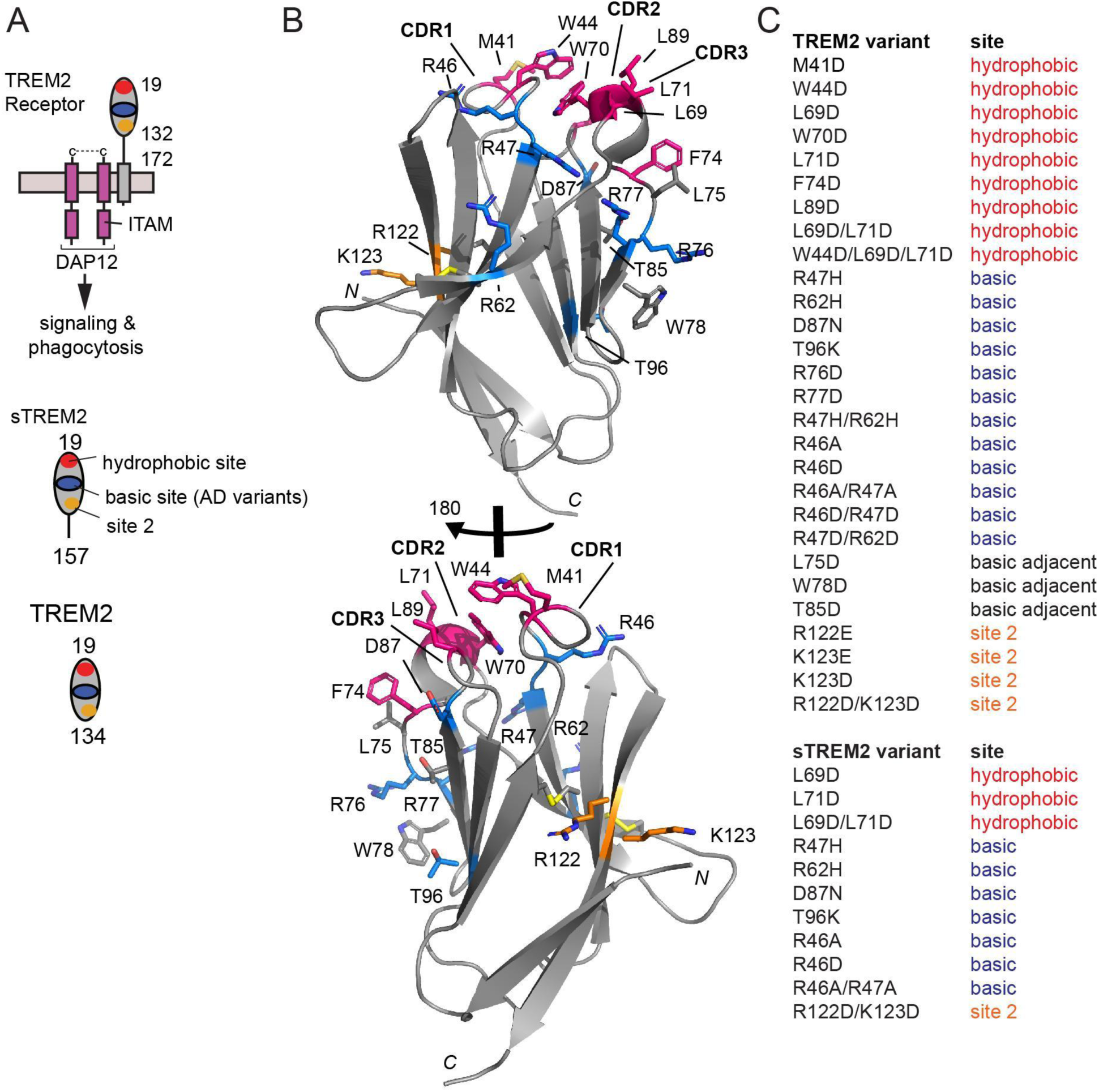
Structure-guided TREM2 variants created to map interactions with ligands. **A)** Schematic of TREM2 receptor on surface of microglia, and TREM2 versions utilized in this manuscript. Cartoon drawing of TREM2 receptor. **B)** Ribbon diagram of TREM2 showing location of residues mutated, colored coded by site (red = hydrophobic; blue = basic; gray = basic adjacent; orange = site 2. **C)** Table of all TREM2 and sTREM2 variants produced for this manuscript or in our previous publications.

In order to understand how TREM2 engages ligands of relevance to AD, we designed a panel of 28 different TREM2 and 11 different sTREM2 variants that could be used in quantitative BLI binding assays to map TREM2 binding surfaces for various ligands. These mutations are mainly located in the previously identified binding surfaces in the hydrophobic site, basic site, and site 2. Here, we used this panel to map binding for the AD-relevant TREM2 ligands apoE, oAβ42, TDP-43, and IL-34. We found that most ligands bound at or near a common site, while IL-34 engaged a unique surface. The results reveal an intriguing promiscuous binding surface on TREM2 and provide information to drive further functional and structural studies, as well as strategies for therapeutically targeting TREM2.

## Methods

### Protein production and purification

TREM2 and sTREM2 variants were cloned into pHLsec vector using the Gibson assembly method. Constructs were verified by sequencing. For TREM2, these constructs were then excised by restriction digest (EcoRI-KpnI), gel purification, and ligation into pHLAvi to encode a C-terminal BirA biotinylation sequence. TREM2 WT and variants were produced in Expi293F cells, purified, and enzymatically biotinylated using BirA as described previously [9, 24]. sTREM2 WT and variants were produced in Expi293F cells, purified from media using NiNTA chromatography, then further purified by gel filtration chromatography on an s200 increase column (Cytiva) in buffer consisting of 20 mM Tris pH 8.5, 150 mM NaCl, and 0.01% azide. ApoE4 was produced as reported previously [9]. Human IL-34 (19-241) was cloned into pHLsec vector [26] and produced in Expi293F cells. The secreted IL-34 was purified from the media using NiNTA chromatography. Full length human TDP-43 was cloned into pET23b to contain a C-terminal 6-His tag. The protein was expressed in *E coli* and purified from insoluble inclusion bodies. Purified TDP-43 was solubilized in a buffer with 6 M guanidine, 10 mM Tris pH 8.0, and 20 mM b-mercaptoethanol. Soluble TDP-43 oligomers were then made 1:100 dilution into 100 mM Tris pH 8.5, 200 mM NaCl, 0.5 M arginine, and 750 mM guanidine. All proteins were analyzed by SDS-PAGE and found to be >95% pure. All proteins were immediately aliquoted, frozen, and stored at −80 deg C for reproducible assays.

### Production of oAβ42 oligomers and biotinylated oAb42 oligomers

WT Ab42 was purchased from Anaspec. Peptides were first dissolved in hexafluoroisopropanol (HFIP) and allowed to air-dry overnight. They were then dissolved in DMSO at a concentration of 100 mM and sonicated for 10 min. These solutions were then diluted into PBS at a concentration of 50 μM and incubated at 4 deg C for 48 hours. Biotinylated oAβ42 was made by mixing N-terminal biotinylated oAβ42 with unlabeled oAβ42 at a 1:20 ratio in DMSO prior to diluting in PBS to form oligomers. Concentrations were then measured by BCA assay, after which the oAβ42 solutions were aliquoted, flash frozen, and stored at −80 C. Freshly thawed aliquots were used for each experiment.

### Binding studies by biolayer interferometry (BLI)

BLI data were collected on an Octet RED384 system (FortéBio). Biotinylated oAβ42 and biotinylated TREM2 were immobilized on streptavidin-coated (SA) biosensors and binding was measured using a running buffer of PBS with 0.1% BSA and 0.005% Tween-20. For experiments with immobilized TREM2 WT and variants, the following proteins were diluted into the running buffer in the following concentration ranges: apoE4 (0.012 - 50 μM), oAβ42 (31.3 - 500 nM), TDP-43 (0.625 - 10 μM), and IL-34 (62.5 - 1000 nM). Data were processed using double-reference subtraction (loaded protein into buffer and biotin-loaded pin into ligand) in ForteBio Data Analysis 9.0.

### BLI competition binding experiments

BLI competition binding experiments were carried out as sequential binding experiments. Biotinylated TREM2 WT was immobilized on SA biosensors then dipped into wells containing either a first ligand or no ligand, then moved to buffer for dissociation, then moved to wells containing a second ligand. The ligand concentrations are listed in the figures and figure legends (**Fig. 5** and **8**).

### Prediction of TREM2-IL-34 binding using computational methods

To identify residues on TREM2 and IL-34 that were likely to involve with protein-protein interactions, we first ran the complete sequence of TREM2 and IL-34 (UniProt accession numbers Q9NZC2 and Q6ZMJ4, respectively) through PredictProtein [27] using ProNA2020 [28], which combines machine learning and homology-based inference to predict if the input sequence is a binding protein and then predict which residues are likely to participate in protein-protein interactions.

Next, we used an in-house hydropathy matching algorithm to predict likely binding sites between TREM2 and IL-34 [9]. We screened the six helices from IL-34 against the TREM2 immunoglobulin domain (residues 18-130). We then screened the three CDR loops that make up the hydrophobic site and the four strands that make up the basic site in TREM2 against the IL-34 sequence (excluding residues 1-20 which make up the signaling peptide). For each screening, the sequence was converted into binary (+ or -) hydrophobicity maps based on the hydrophobicity sign of each residue by the Kyte and Doolittle scale [29]. The hydrophobicity maps for each potential binding motif were screened in both forward and reverse orientations against the entire hydrophobicity map of the other protein. The percent match was calculated as the percentage of dividing complementary (+/-) pairs to all matched pairs. The degree of complementary hydropathy (Eq. 1[30]) was calculated based on the Kyte Doolittle hydropathy index.

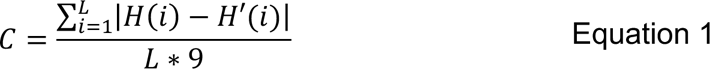

In Eq. 1, is the degree of complementary hydropathy, and are the hydropathy indices of the residues in the motif and target sequences respectively at position, and is the length of the motif. The degree of complementary hydropathy can range from 0 to 1. A predicted hit was considered good if the percent match was greater than 75% and the degree of complementary hydropathy was greater than 0.5.

To further predict the interactions between TREM2 and IL-34, we implemented protein-protein docking using HADDOCK [31] with TREM2 residues of R47, R62, L75, R76, and R77 that could cause at least a 2-fold decrease in K_D_ when mutated verse WT, as restraints. IL-34 domains that were predicted as potential binding sites for TREM2 from the hydropathy screening were set as the preferred binding domains for TREM2 in protein-protein docking. Prior to docking, we energy minimized the IL-34 structure (PDBID: 4DKC [32] in Amber18 and used a previously equilibrated TREM2 structure (PDBID: 5UD7[25]). The hydrogen bond analysis and electrostatic surface potential analysis of the top predicted complex from HADDOCK was analyzed in PyMOL.

## Results

### Creation of a TREM2 variant protein library to comprehensively evaluate TREM2-ligand interactions

In order to comprehensively investigate the involvement of putative TREM2 binding surfaces in engaging various ligands, we designed a structure-guided library of TREM2 and sTREM2 variants (**Fig 1**). Due to the anionic or low pI nature of most TREM2 ligands, most variants were designed at mutations to aspartate (D) and most were point variants, with the exception of some double and triple mutants designed at each site (hydrophobic, basic, site 2). In total, we have created 28 TREM2 variants and 11 sTREM2 variants (**Fig 1**). All TREM2 variants were cloned into a vector that contained a specific biotinylation sequence at the C-terminus. Thus when immobilized on streptavidin-coated BLI pins, these proteins are presented to ligands in the same orientation and oligomerization state as they are on the cell surface. All proteins were expressed in mammalian cells as we previously published [33], thus they contain similar post translational modifications (glycosylation and disulfide bonds) to the native proteins expressed by microglia. The purified biotinylated TREM2 and sTREM2 proteins were aliquoted and frozen, allowing for reproducibility. This protein library represents a powerful tool for mapping TREM2 interactions with ligands.

### Mutations to the TREM2 hydrophobic site can ablate binding to apoE4

In a previous study, we showed that apoE4 bound to TREM2 with K_d_ = 281 nM, and that AD risk variants in TREM2 (R47H, R62H, and T96K) did not grossly impact binding [9]. Instead, we found that point mutations to the hydrophobic site, namely that mutants L69D and W70D, decreased binding affinity by nearly 200-fold (**Fig. 2B** and **Table 1**). In order to comprehensively extend these studies, we carried out further BLI experiments to complete characterizing the most common AD risk variants, and used TREM2 mutants at all three binding sites. Surprisingly, while the TREM2 AD variants R47H, R62H, and T96K had previously shown small decreases in binding affinity for apoE4 (2-3 fold decrease in K_D_, **Table 1**), we found that AD risk variant D87N showed a dramatic 11-fold increase in affinity for apoE4 (Kd = 25 nM; **Fig 2F** and **Table 1**). Next we evaluated the impact of double and triple mutations to the three putative binding surfaces on TREM2. Strikingly, we found that double (L69D/L71D) and triple (W44D/L69D/L71D) mutations at the hydrophobic site completely ablated binding to apoE4 (**Fig 2D,E** and **Table 1**). In stark contrast, double mutations to the basic site (R46A/R47A) and site 2 (R122D/K123D) did not did not largely impact binding to apoE4 (**Fig. 2C & I, Table 1**). Two point mutations in the basic site, R76D and R77D, did display 5-fold decreases in K_D_ for apoE4 (**Fig 2G,H**. **Table 1**), suggesting that these residues might be partially involved in engaging apoE4. However, these residues are directly adjacent to the hydrophobic site, and might impact conformation of CDR2. In summary, our results suggest that apoE4 primarily engages the hydrophobic site on TREM2.

**Figure 2.**
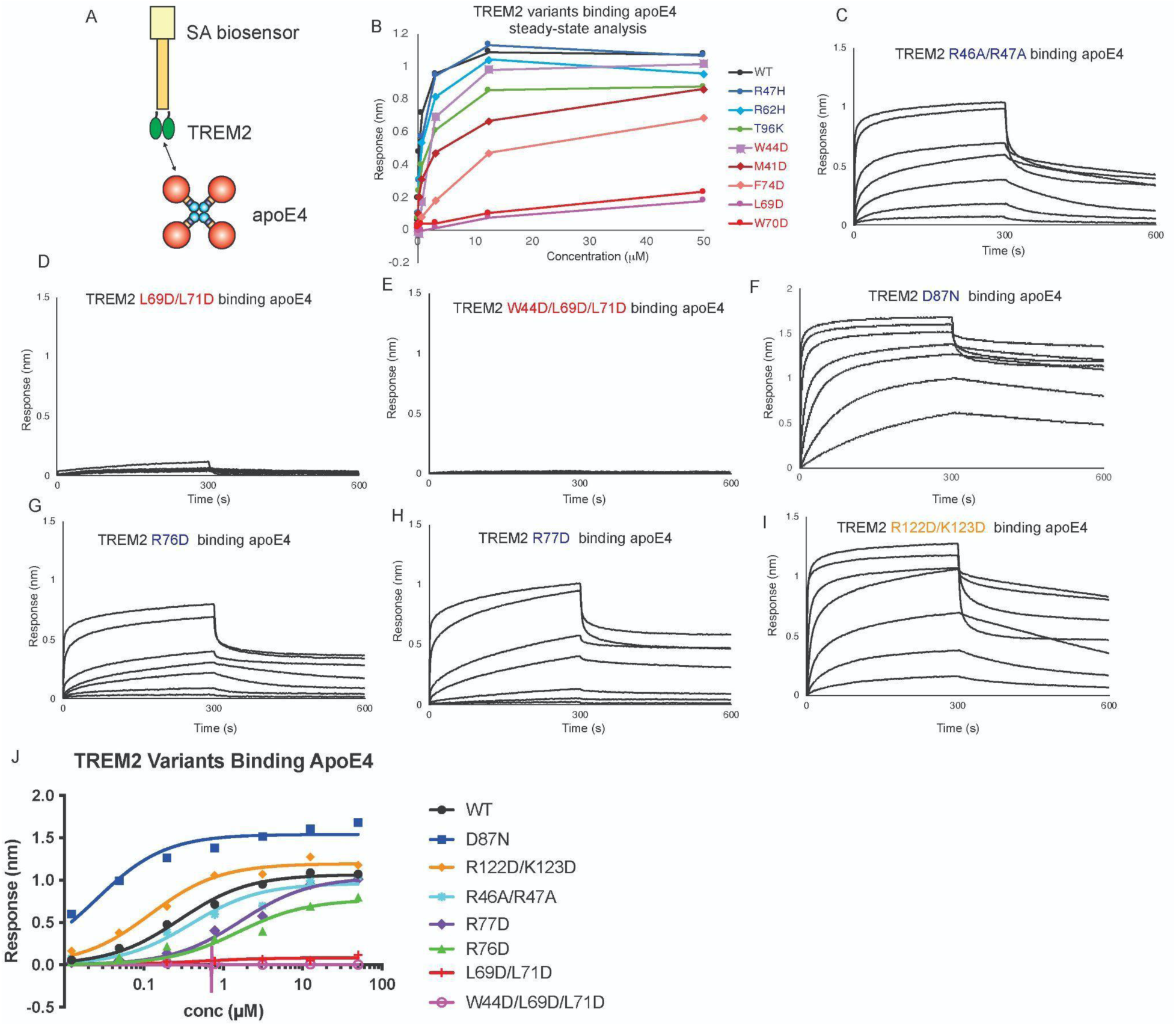
Mutations to TREM2 hydrophobic site ablate binding to apoE4. Immobilized TREM2 WT and variants were probed for binding to apoE4 (0.012 - 50 microM). **(A)** Scheme of experiment. **B)** Summary of steady-state binding for TREM2 WT and variants from our previous publication (REF). **C-J)** BLI sensorgrams for TREM2 R46A/R47A, **D)** L69D/L71D, **E)** W44D/L69D/L71D, **F)** D87N, **G)** R76D, **H)** R77D, **I)** R122D/K123D binding to apoE4 (0.012 - 50 microM). Double-reference subtracted data shown in black. **K)** Steady state analysis and non-linear fits to derive K_D_ from data shown in **C-J**. The derived K_D_s are listed in **Table 1**.

**Table 1.**
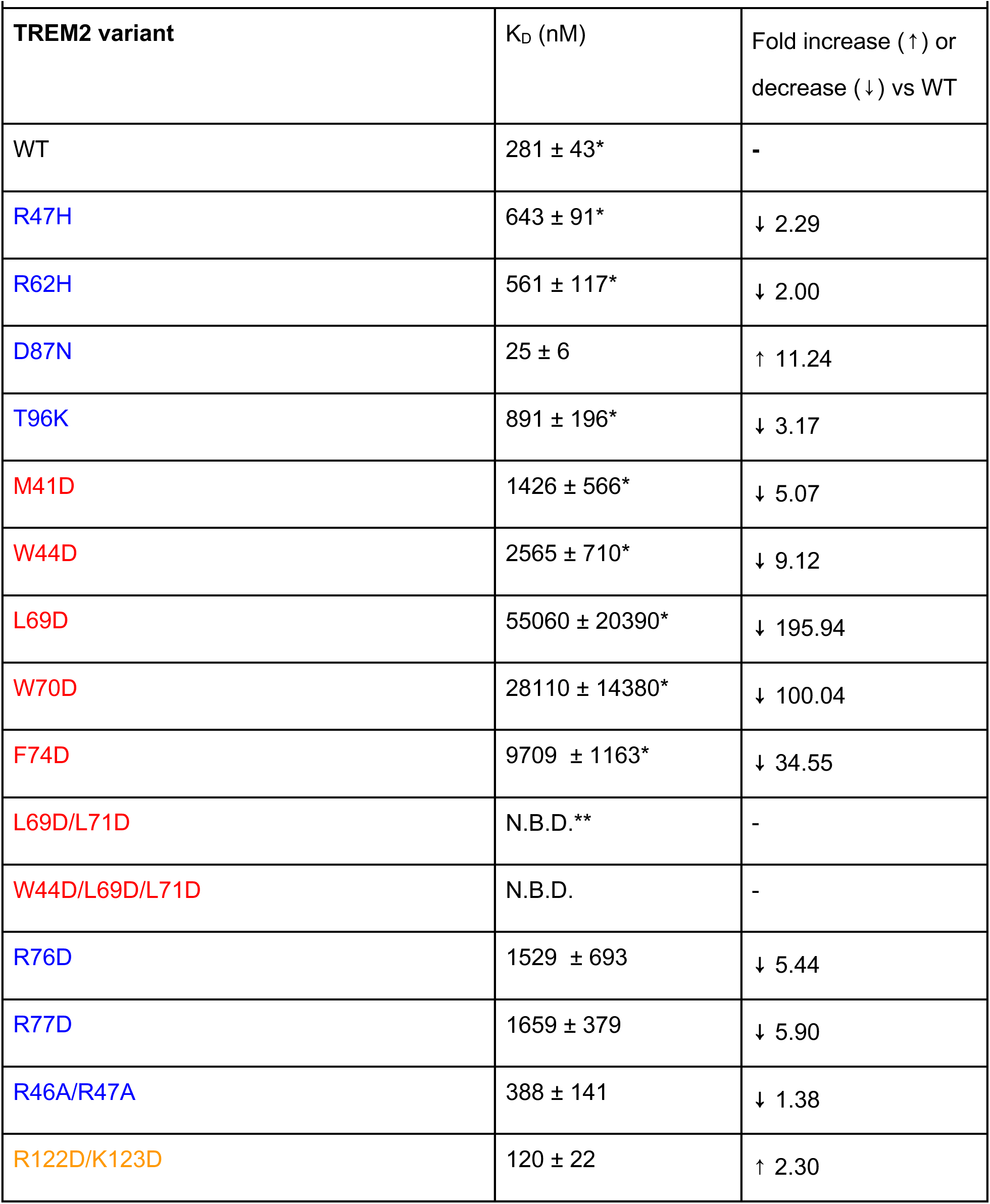
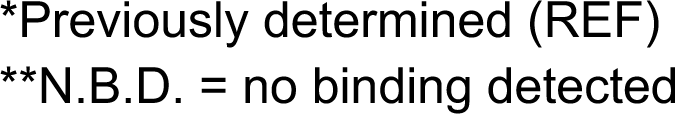
TREM2 variants binding to apoE4.

| Table 1. TREM2 variants binding to apoE4 |  |  |
| --- | --- | --- |
| TREM2 variant | K<sub>D</sub> (nM) | Fold increase (↑) or decrease (↓) vs WT |
| WT | 281 ± 43* | - |
| R47H | 643 ± 91* | ↓ 2.29 |
| R62H | 561 ± 117* | ↓ 2.00 |
| D87N | 25 ± 6 | ↑ 11.24 |
| T96K | 891 ± 196* | ↓ 3.17 |
| M41D | 1426 ± 566* | ↓ 5.07 |
| W44D | 2565 ± 710* | ↓ 9.12 |
| L69D | 55060 ± 20390* | ↓ 195.94 |
| W70D | 28110 ± 14380* | ↓ 100.04 |
| F74D | 9709 ± 1163* | ↓ 34.55 |
| L69D/L71D | N.B.D.** | - |
| W44D/L69D/L71D | N.B.D. | - |
| R76D | 1529 ± 693 | ↓ 5.44 |
| R77D | 1659 ± 379 | ↓ 5.90 |
| R46A/R47A | 388 ± 141 | ↓ 1.38 |
| R122D/K123D | 120 ± 22 | ↑ 2.30 |

\*Previously determined (REF)
\*\*N.B.D. = no binding detected

### Mutations to the TREM2 hydrophobic site ablate binding to oAβ42

In accompanying work, we determined the structural and functional determinants for TREM2 engaging oAβ42. In this work, we show that oAβ42 engages the hydrophobic site on TREM2 utilizing the N-terminal portion of the Aβ42 peptide. Double and triple mutations to the hydrophobic site completely ablated binding to oAβ42. Given the oligomeric nature of oAβ42, we wanted to comprehensively examine whether other surfaces on TREM2 might be partially involved in engaging oAβ42. To do this, we probed the binding of oAβ42 to TREM2 using a variety of TREM2 variants that contained single or multiple mutations at each of the three putative binding sites on TREM2. With TREM2 immobilized on the pin, oAβ42 bound with high affinity (Kd = 42.5 nM) (**Fig. 3A,B**). Because oAβ42 does not noticeably dissociate from immobilized TREM2 in these BLI experiments, titrations were carried out in a vertical orientation with each pin used to probe a single concentration of oAβ42. Care was taken to ensure that pins were loaded with nearly identical amounts of TREM2 proteins so that magnitudes could be compared. For the TREM2 hydrophobic site variants, we found that point mutations decreased binding, while double (L69D/L71D) and triple (W44D/L69D/L71D)0 mutations completely ablated binding **(Fig. 3L**). We found that point mutations to the basic site (R76D, R77D) showed slightly decreased binding (**Fig 3C, D**). The basic site AD variant R62H only slightly decreased binding to oAβ42 (**Fig. 3J**), while D87N showed binding near that of WT TREM2 (**FIg. 3K**). A basic site double mutant, R47H/R62H, showed slightly decreased binding, but was similar to the R62H variant (**Fig. 3E, M**). Point mutations to site 2 (R122E, K123D, K123E) also showed slightly decreased binding to oAβ42 (**Fig. 3F,G,H**) as did the site 2 double mutant R122D/K123D **(Fig. 3I, N**). In summary, mutations to the hydrophobic site reduced or ablated association while mutations to the basic site and site 2 either did not impact or slightly lowered association.

**Figure 3.**
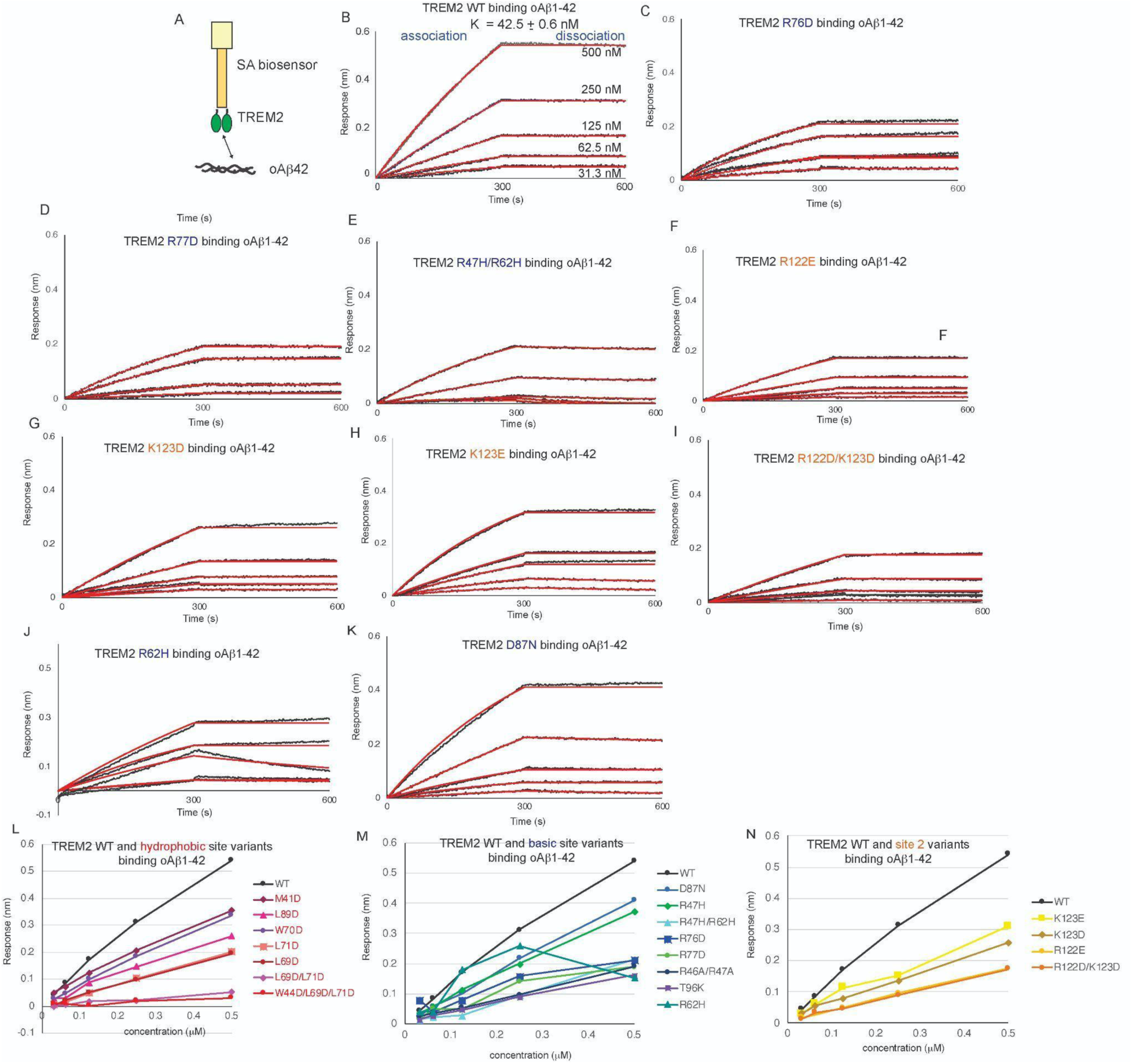
Mutations to TREM2 hydrophobic site disrupt binding to oligomeric Aβ42. Immobilized TREM2 WT and variants were probed for binding to oAβ42 (31.3 - 500 nM). **(A)** Scheme of experiment. B-I) BLI response for TREM2 **B)** WT, **C)** R76D, **D)** R77D, **E)** R47H/R62H, **F)** R122E, **G)** K123D, **H)** K123E, **I)** R122D/K123D, **J)** R62H. **K)** D87N binding to oAb42 (31.3 - 500 nM). Double-reference subtracted data (black) overlayed with 1:1 kinetic fits (red). K_D_ derived from kinetic fits. Data representative of at least two independent experiments. **L-N)** Summary of BLI maximum binding response versus concentration for oAβ42 binding to TREM2 variants grouped by **L)** hydrophobic site, **M)** basic site, and **N)** site 2.

Given the oligomeric nature of oAβ42, it is likely that a single oligomer could engage multiple TREM2 molecules immobilized on the pin, leading to enhanced binding due to avidity affects. In order to more accurately assess the engagement of a single TREM2 molecule with oAβ42, we carried out experiments in the opposite orientation with biotinylated oAβ42 immobilized on the pin and sTREM2 WT and variants in the well (**Fig 4A**). In the absence of avidity, single site mutations at critical residues should have a more pronounced impact on binding. In this orientation, single point mutations to the hydrophobic site (L69D, W70D) ablated binding to oAβ42 as did the double mutation L69D/L71D (**Fig 4C**). In contrast, basic site point mutations (R46A, R47D, R62H, R47H, D87N, T96K) showed minor reductions in binding to oAβ42, and double mutation to the basic site (R46A/R47A) also did not greatly reduce binding compared to WT sTREM2. (**Fig. 4C-F**). Kinetic analysis indicated basic site mutations reduced K_d1_ by about 2 orders of magnitude (Table 2), as did a double mutation at site 2 (R122D/K123D) (**Fig 4C, G**). Overall, however, these decreases were minor compared to the complete ablation caused by point mutations in the hydrophobic site. Taken altogether, these comprehensive binding studies indicate that the TREM2 hydrophobic site is the primary binding site for oAβ42, while other surfaces of TREM2 might be involved in some extended contacts.

**Figure 4.**
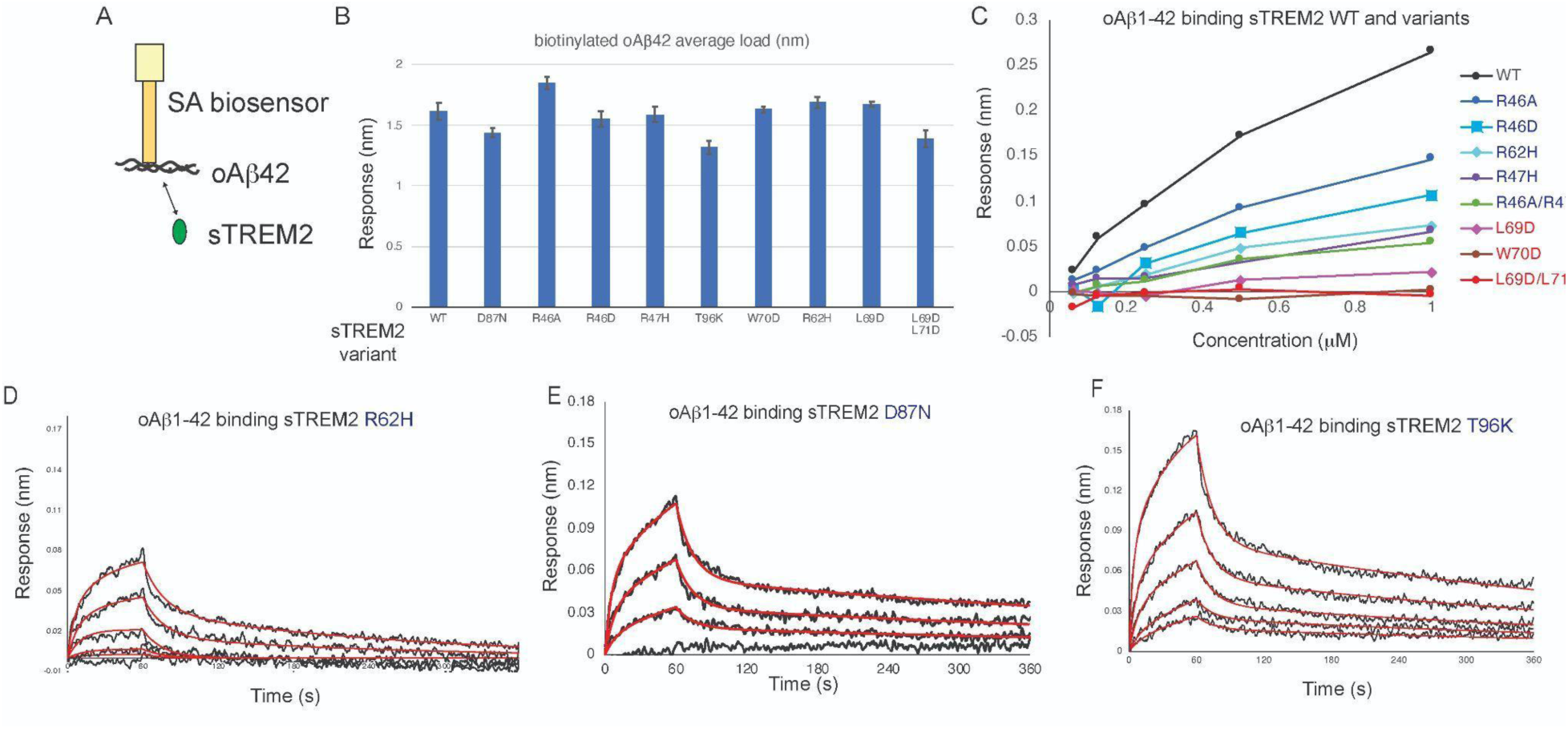
Mutations to the hydrophobic site prevent sTREM2 binding to oAβ42, mutations to the basic site do not. **A)** Scheme of experiment. **B)** Amount of biotinylated oAβ42 immobilized in experiments with sTREM2 variants. Values shown are the average of five loads, with standard errors shown. C) Summary of maximum response binding for TREM2 WT and variants from our accompanying publication. D-F) BLI response for sTREM2 **D)** R62H, **E)** D87N, **F)** T96K binding to oAβ42. sTREM2 concentration range was 0.625 - 1.0 μM. Double-reference subtracted data (black) overlayed with 2:1 kinetic fits (red). Data representative of at least two independent experiments.

**Figure 5.**
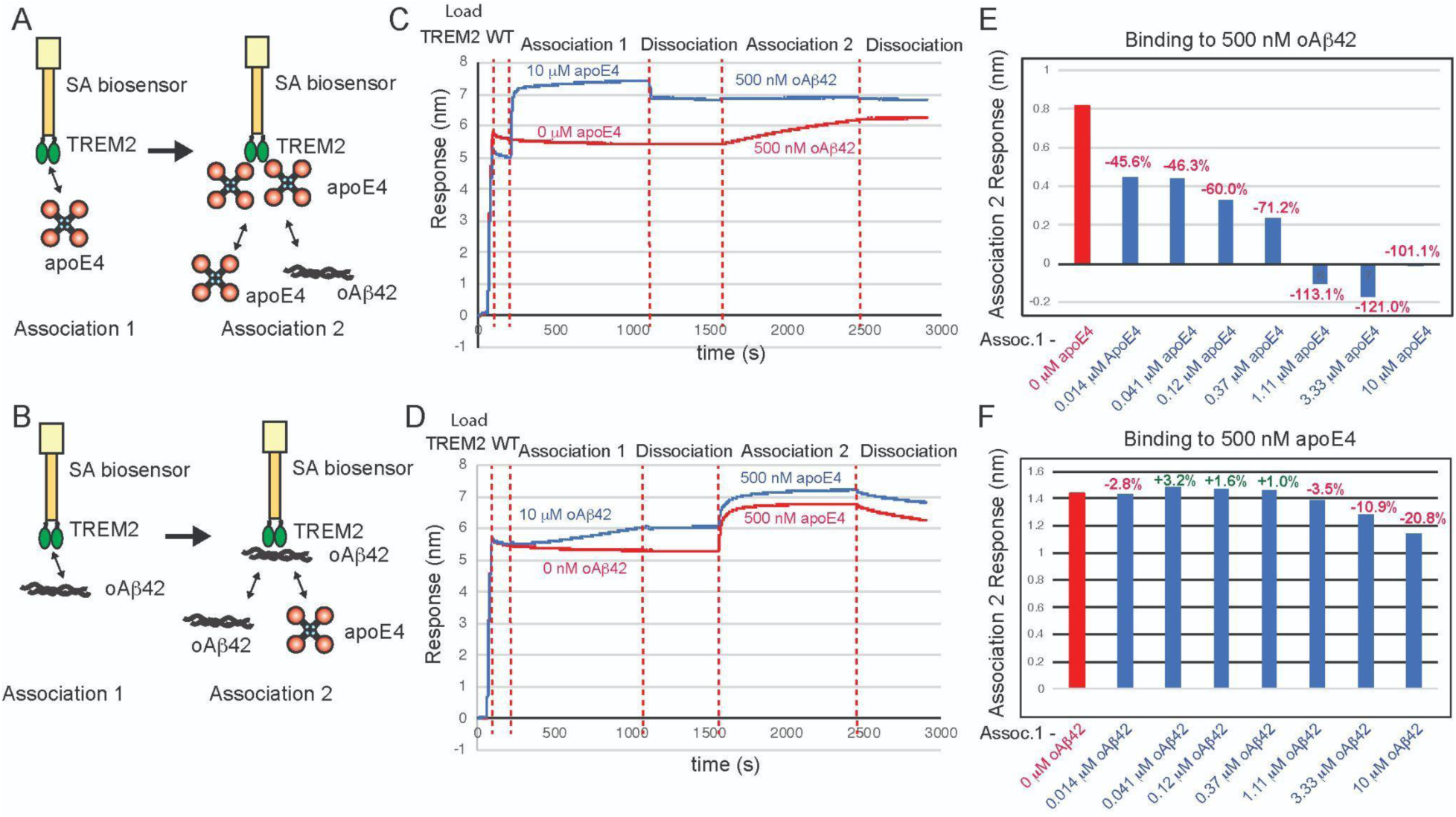
ApoE4 and oAβ42 compete for binding to TREM2. **A & B)** Schematic of competition binding BLI experiments. **C&D)** BLI sensograms for **C)** apoE4 competing oAβ42 binding to TREM2 and **D)** oAβ42 competing apoE4 binding to TREM2. Red sensorgrams are TREM2 binding to **C)** 500 nM oAβ42 or **D)** 500 nM apoE4 alone while blue sensorgrams show competition experiments where **C)** 10 μM apoE4 or **D)** 10 μM oAβ42 are bound first. **E&F)** BLI binding magnitudes for TREM2 binding to **E)** 500 nM oAβ42 when pre-binding increasing concentrations of apoE4 or **F)** 500 nM apoE4 when pre-binding increasing concentrations of oAβ42.

**Table 2:** sTREM2 binding kinetics to oAB42.

| Table 2: sTREM2 binding kinetics to oAB42 |  |  |
| --- | --- | --- |
| sTREM2 variant | Kd<sub>1</sub> (M) | Kd<sub>2</sub> (M) |
| WT | 6.30E-09 | 4.78E-07 |
| L69D | NBD | NBD |
| W70D | NBD | NBD |
| L69D/L71D | NBD | NBD |
| R47H |  |  |
| R62H | 1.87E-07 | 5.11E-07 |
| D87N | 1.85E-07 | 3.89E-07 |
| T96K | 9.31E-08 | 3.83E-07 |
| R46A | 3.20E-09 | 3.39E-04 |
| R46D | 1.40E-07 | 5.79E-07 |
| R46A/R47A | 2.243E-08 | 6.308E-07 |
| R122D/K123D | 1.912E-07 | 1.835E-05 |

### Aβ42 and apoE compete for binding to the TREM2

Since our mapping studies indicated that both apoE4 and oAβ42 bind to the hydrophobic site on TREM2, we conducted assays to examine if they compete for a common binding site. In these competition binding assays, TREM2 was immobilized on the pin and then allowed to associate with titrating concentrations of apoE4, then dipped into wells containing 500 nM oAβ42 (**Fig 5A**). We found that apoE4 robustly inhibited the binding of oAβ42 to TREM2, with concentrations of apoE4 above 1000 nM able to completely block binding of oAβ42 to TREM2 (**Fig 5C,E**). We then carried out the experiment in the opposite order, with TREM2 first binding to titrating concentrations of oAβ42 (**Fig. 5B**). In this orientation, we found that oAβ42 only weakly inhibited binding of apoE4 to TREM2, with modest inhibition noted at concentrations above 1000 nM (**Fig. 5D,F**). These results indicate that apoE4 and oAβ42 share an overlapping binding site on TREM2, and demonstrate that aopE4 can more strongly compete for binding to TREM2 in the presence of oAβ42.

### Mutations to the TREM2 hydrophobic site severely inhibit binding to TDP-43

TDP-43 aggregate accumulations are found in most ALS patients, and are also found in individuals with FTD and AD. TREM2 was recently identified as a receptor for TDP-43 oligomers, with engagement triggering phagocytic clearance of TDP-43 oligomers by microglia [19]. In that report, direct interaction between TREM2 and soluble TDP-43 oligomers was demonstrated using SPR. In order to identify the binding site on TREM2 for oligomeric TDP-43, we prepared TDP-43 oligomers in a similar manner and used BLI to probe interactions with TREM2 variants across the three TREM2 binding sites. With TREM2 immobilized on the pin, we found that it bound TDP-43 robustly (**Figure 6A, B)**. Due to the oligomeric nature of TDP-43, binding curves were biphasic, so 1:1 kinetic fits were not appropriate; therefore, binding curves were only qualitatively evaluated. Mutations to the basic site (R47H, R62H, D87N) and site 2 resulted in binding magnitudes comparable to WT (**Fig 6 G, H, I, K**). A double mutant at the basic site (R46A/R47D) displayed slightly reduced binding (**Fig 6C, K**). Similarly, double mutation R122D/K123D at site 2 resulted in slightly increased binding as compared to WT (**Figure 6J,K**). In stark contrast, single (W70D), double (L69D/L71D), and triple mutations (W44D/L69D/L71D) to the hydrophobic site completely ablated binding to TDP-43 (**Table 4**, **Figure 6D,E,F, K**). Altogether, these results suggest that TDP-43 oligomers engage TREM2 at the hydrophobic site near residue 70.

**Figure 6.**
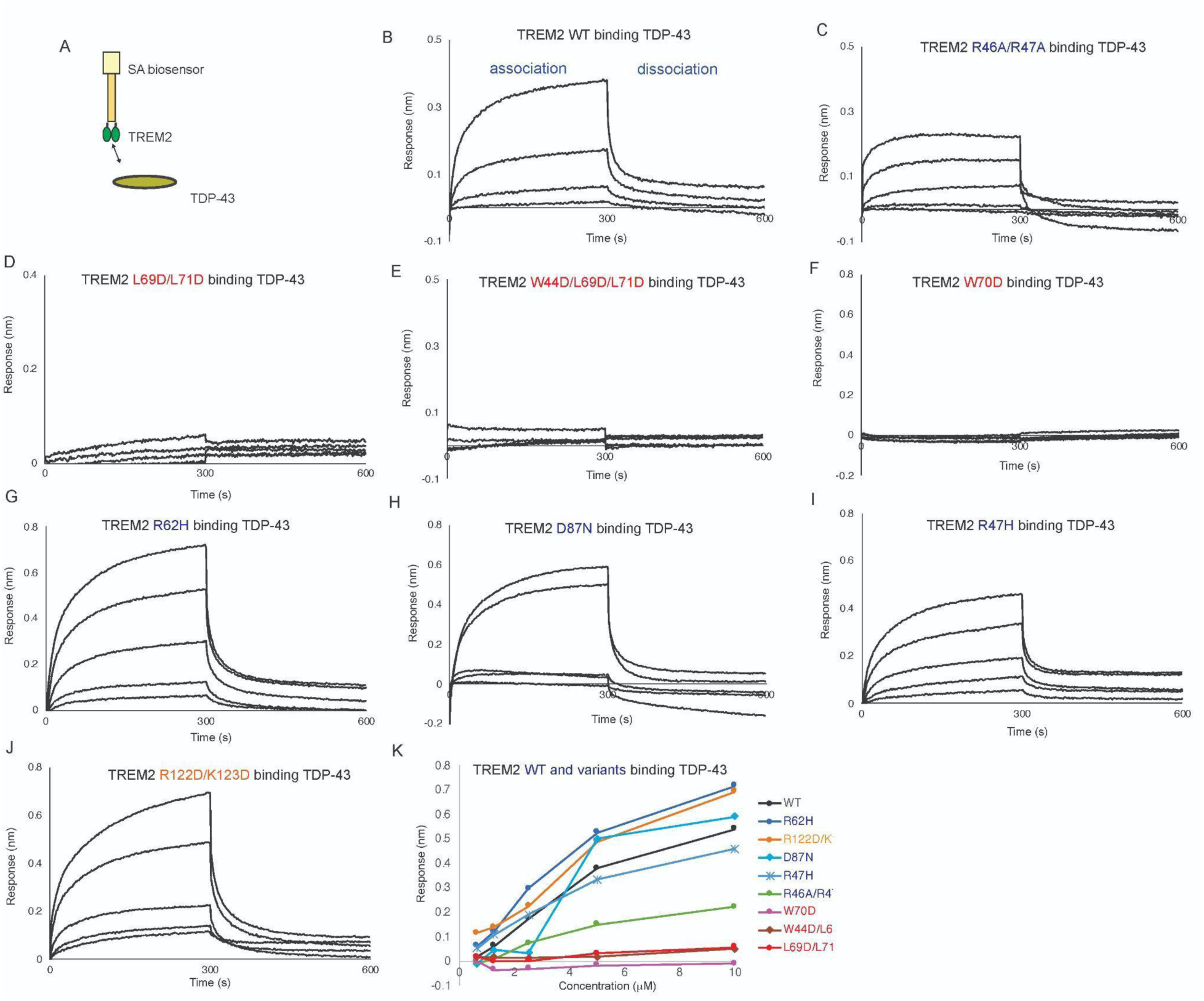
Mutations to TREM2 hydrophobic site ablate binding to TDP-43. Immobilized TREM2 WT and variants were probed for binding to TDP-43 (0.625 – 10 μM). **(A)** Scheme of experiment. **B-F)** BLI response for TREM2 **B)** WT, **C)** R46A/R47A, L69D/L71D, **E)** W44D/L69D/L71D, **F)** W70D**, G)** R62H, **H)** D87N, **I)** R47H, **J)** R122D/K123D binding to TDP-43 (0.625 - 10 μM). Double-reference subtracted data (black) is shown. **K)** Summary of BLI steady-state binding response versus concentration for TDP-43 binding to TREM2 variants.

### Mutations to the TREM2 basic site around R76 severely inhibit binding to IL-34

IL-34 was recently identified as a signaling ligand for TREM2 [22] In order to map the binding surface for IL-34 on TREM2, we carried out BLI binding studies with our TREM2 variant library. With TREM2 WT immobilized on the BLI pin, we found that IL-34 bound with high affinity (K_d_ = 16.5 nM) (**Fig. 7A,B** and **Table 3**). In contrast to the other ligands studied here, IL-34 bound to some TREM2 AD risk variants (R47H, R62H) with slightly lower affinity, showing around a 4-fold decrease in K_D_ (**Fig. 7C Table 3**). The AD risk variants D87N and T96K did not largely impact IL-34 binding (**Fig. 7D,E and Table 3**). We further probed the basic site and found that the R77D variant showed a nearly 7-fold decrease in affinity (K_D_ = 114 nM, **Fig 7J** and **Table 3**) while the R76D mutant displayed no binding to IL-34 at the concentration range probed (**Fig 7I** and **Table 3**). We further probed this region and introduced mutations at residues adjacent to R76. These variants (L75D, W78D, T85D) did not impact binding to IL-34 (**Figure 7L-N** and **Table 3**). Another mutation to the basic site, R46D, also did not impact binding to IL-34 (**Fig 7F** and **Table 3**). We then probed mutations at the hydrophobic site and site 2. The site 2 variant (R122E) did not impact binding (**Fig 7O**), nor did the site 2 double mutant R122D/K123D (**Fig. 7P**). Most notably, the double and triple hydrophobic site variants L69D/L71D (**Fig. 7H**) and W44D/L69D/L71D (**Fig. 7G**) did not affect binding to IL-34. These results suggest that IL-34 binds to the TREM2 basic site in a region centered on R76.

**Figure 7.**
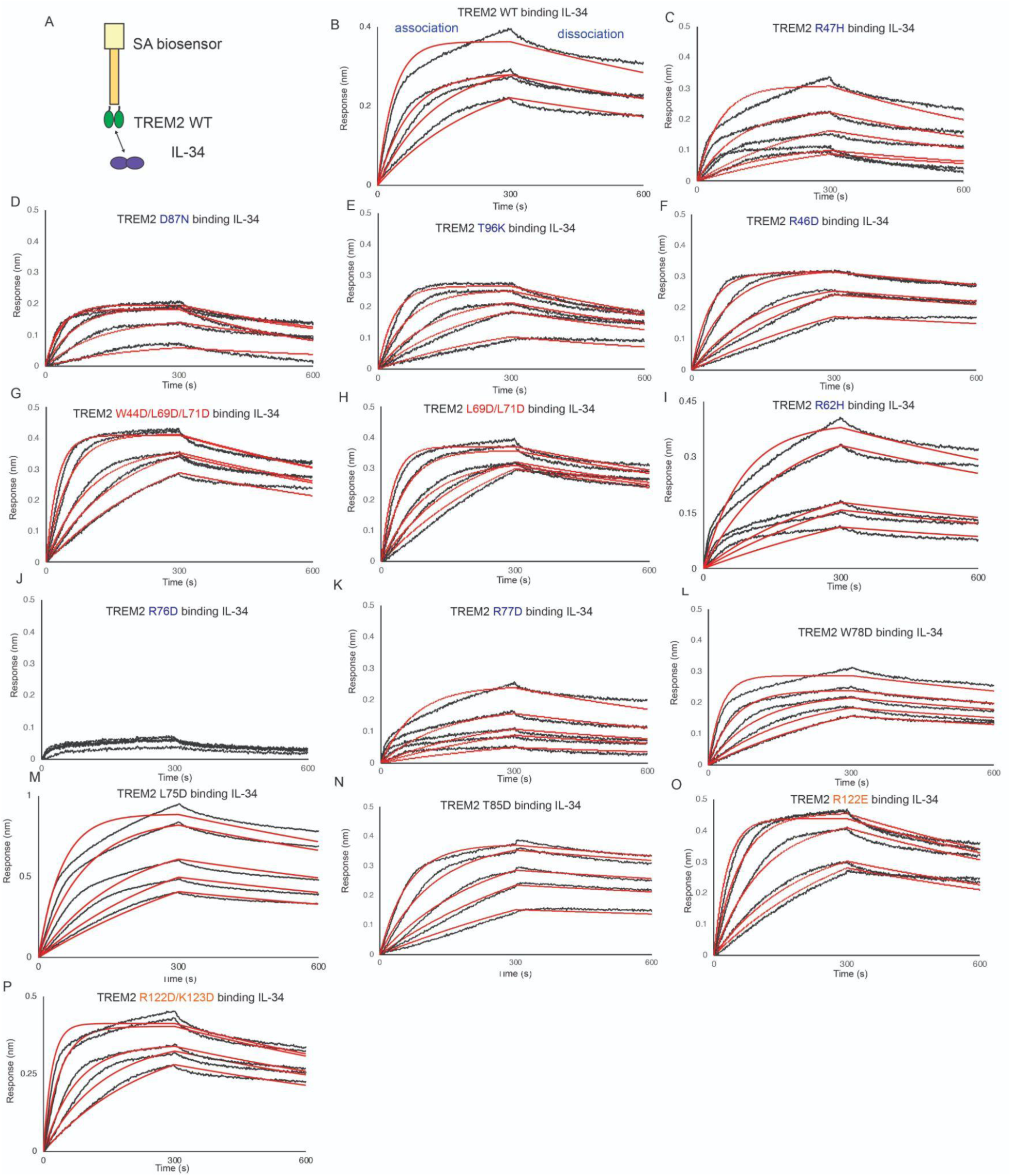
TREM2 basic site variants centered around R76 inhibit binding to IL-34. Immobilized TREM2 was probed for binding to IL-34 (62.5 - 1000 nM). **(A)** Scheme of experiment. **(B-N)**BLI sensorgrams for IL-34 binding to TREM2 **(B)** WT, **(C)** R47H, **(D)** D87N, **(E)** T96K, **(F)** R46D, **(G)** W44D/L69D/L71D, **(H)** L69D/L71D, **(I)** R62H, **(J)** R76D, **(K)** R77D, **(L)** W78D, **(M)** L75D, **(N)** T85D, **(O)** R122E, (**P**) R122D/K123D. Black = BLI sensorgrams; red = 1:1 kinetic fits. Results shown in **Table 3**.

**Figure 8.**
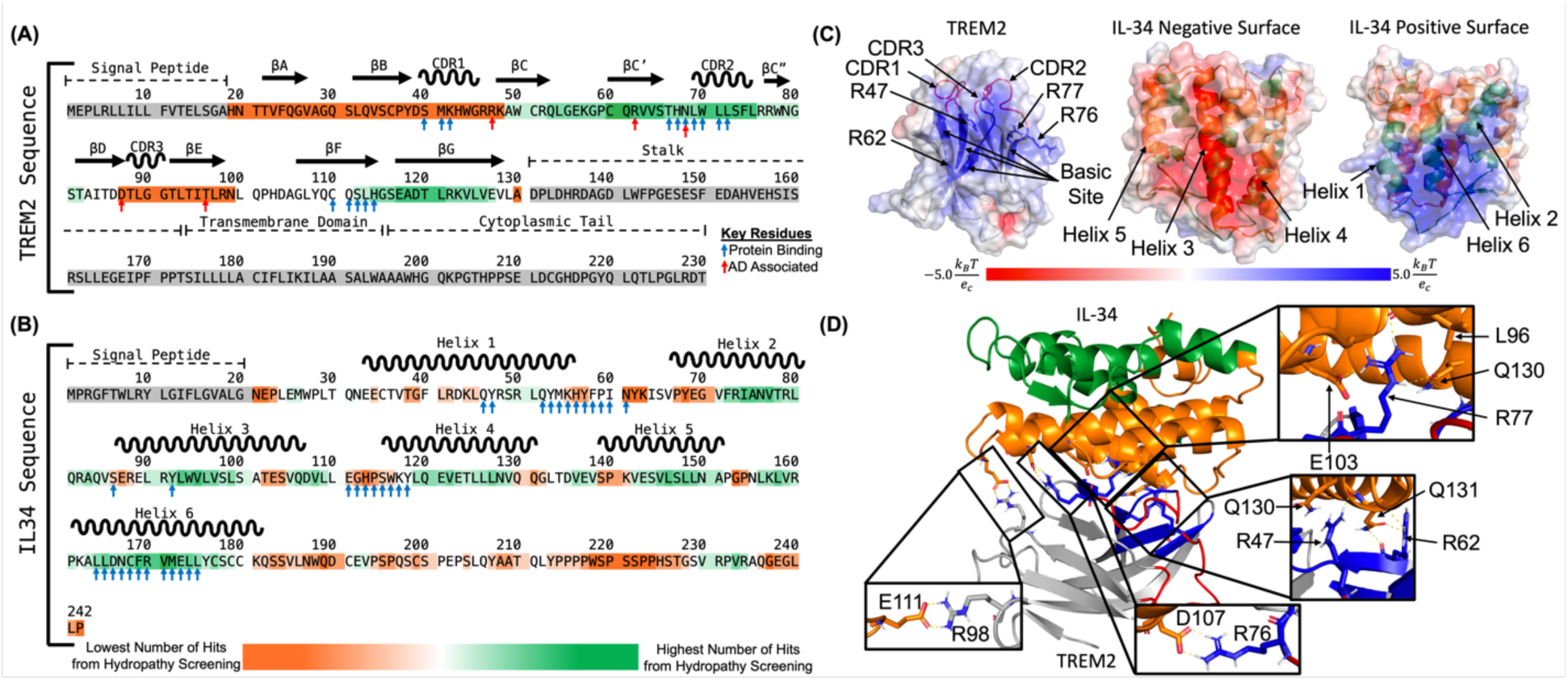
Computational prediction of the interactions between TREM2 basic site and IL-34 negatively charged surface. **(A)** The complete sequence of human TREM2 showing predicted potential key residues and binding regions between residues 49-82 and 112-127 (Basic site and CDR2) for IL-34. Blue arrows denote residues predicted to be important for protein binding via PredictProtein and red arrows denote residues that increase Alzheimer’s disease risk. Residues in TREM2 immunoglobulin domain are highlighted ranging from orange to green. Residues highlighted in the darkest green had the highest number of hits and residues highlighted in the darkest orange had the lowest number of hits. Residues highlighted in grey were not screened. **(B)** The complete sequence of IL-34 showing predicted potential key residues and binding regions between residues 71-85, 90-100, 119-129, 142-151, and 156-179 for TREM2. Blue arrows denote residues predicted to be important for protein binding via PredictProtein. Residues highlighted in the darkest green had the highest number of hits and residues highlighted in the darkest orange had the lowest number of hits. Residues highlighted in grey were not screened. **(C)** Electrostatic surface potential maps showing positively charged TREM2 basic site, negatively charged IL34 surface (made up of helices 3, 4, and 5), and positively charged IL34 surface (made up of helices 1, 2, and 6). Regions with positive electrostatic surface potential are shown as blue, regions with negative electrostatic surface potential are shown as red, and neutral regions are shown as white. TREM2 is shown as grey with the basic site show as blue cartoon and the hydrophobic site shown as red cartoon. Key regions and residues for binding are labeled. IL-34 is shown as cartoon with residues making up the negatively charged surface colored in orange and residues making up the positively charged surface colored in green. The six helices are labeled. **(D)** Predicted complex structure of TREM2 with IL-34 shows the negatively charged surface of IL-34 (Helices 3, 4, and 5) interaction with the positively charged basic site of TREM2, and key TREM2 residues identified through BLI forming hydrogen bonds and salt bridges with IL-34 residues. TREM2 is shown in grey, TREM2 basic site is shown as blue, and TREM2 hydrophobic site is shown as red. Residues making up IL-34 negatively charged surface are shown as orange and residues making up IL-34 positively charged surface are shown as green. Hydrogen bonds and salt bridges between key identified TREM2 residues from BLI and residues from IL-34 are shown as sticks and the interactions are shown as dashed yellow lines.

**Table 3.** TREM2 variants binding to IL-34.

| Table 3. TREM2 variants binding to IL-34 |  |  |
| --- | --- | --- |
| TREM2 variant | K<sub>D</sub> (nM) | Fold decrease vs WT |
| WT | 16.5 ± 0.2 | - |
| R47H | 79.8 ± 2.4 | ↓4.8 |
| R62H | 75 | ↓4.5 |
| D87N | 28.4 ± 0.5 | ↓1.7 |
| T96K | 23.7 ± 0.4 | ↓1.4 |
| R46D | 18.0 ± 0.3 | ↓1.1 |
| R76D | N.B.D | - |
| R77D | 114 ± 4 | ↓6.9 |
| L75D | 36.6 ± 0.9 | ↓2.2 |
| W78D | 17.2 ± 0.4 | ↓1.04 |
| T85D | 21.5 ± 0.6 | ↓1.3 |
| R122E | 29.4 ± 0.4 | ↓1.8 |
| R122D/K123D | 18.5 ± 0.3 | ↓1.1 |
| L69D/L71D | 20.2 ± 0.3 | ↓1.2 |
| W44D/L69D/L71D | 25.8 ± 0.3 | ↓1.6 |

### A predicted structure for the TREM2/IL-34 complex

A recent manuscript presented a computational prediction for the TREM2/IL-34 complex [22]. In this report, IL-34 was predicted to bind TREM2 at the hydrophobic site, with contact residues including W44, W70, and L71. However, our experimental results show that mutations to the hydrophobic site do not impact binding to TREM2, and instead show that the IL-34 binding site is on the basic site, centered on R76. Therefore, we further undertook computational prediction of the binding sites between TREM2 and IL- 34 to support our experimental data.

We first identified key residues of TREM2 and IL-34 that were most likely involved in protein-protein interactions using a machine learning and homology-based inference approach [27]. In TREM2, 14 residues were identified as likely involved in binding. These include residues in CDR1 (residues 40-42) and CDR2 (residues 69-72) as well as basic site residues (residues 66-68 and 112-114) (**Fig. 8A**). The residues predicted around CDR2 are directly surrounded by residues that can strongly inhibit IL-34 binding when mutated (**Table 3**) including R62, R76, and R77. Both residues 112-114 and residues 66-68 are spatially adjacent to another residue, R47, that can strongly inhibit IL-34 binding when mutated. In IL-34 there were 34 residues identified as likely involved in protein-protein binding. These residues existed primarily at the end of Helix 1, the beginning of Helix 4, and throughout Helix 6, suggesting the binding likely occurs in the helix bundle (**Fig. 8B**).

To further identify regions of TREM2 and IL-34 that could be responsible for binding, we used a sequence-based hydropathy mapping approach. The structure of IL-34 can be broken into six helices. Each helix sequence was screened against the sequence of TREM2 immunoglobulin (Ig) domain to identify potential regions of binding. Scanning both the forward and reverse residue sequences of the six helices, hits with more than 75% percent match and greater than 0.5 degree of complementary hydropathy were considered successful. Based on our criteria, helices 2, 5, and 6 all had good hits targeting a combination of TREM2 basic site and TREM2 CDR2 (**Table S1**). When the successful hits were clustered on the TREM2 sequence, we found two regions for predicted binding, residues 49-82 and residues 112-127 (**Fig 8A**). These two strands contain most of the residues in the basic site, as well as the entirety of CDR2, consistent with the BLI results (**Table 3**). Similarly, we screened the three TREM2 CDR loops (hydrophobic regions – CDR1: 39-46, CDR2: 69-75, CDR3: 88-91) as well as the four TREM2 strands that make up the basic site (residues 47-50, 62-68, 76-78, and 112-114) separately across the sequence of IL-34 to identify potential binding regions between TREM2 and IL-34. We scanned both the forward and reverse residue sequences of the three CDR loops and four basic site strands in TREM2 against the sequence of IL-34, where hits with more than 75% percent match and greater than 0.5 degree of complementary hydropathy were considered successful. From these results, we noted the largest amount of good hits came from the basic site residues, as well as CDR2 in TREM2 (**Table S2**). This again matches well with our predicted binding regions on TREM2, as well as the BLI results, which showed mutations of the residues from TREM2 basic site were able to strongly disrupt TREM2/IL-34 binding (**Table 3**). We clustered the good hits on the sequence of IL-34 and noted five regions of potential interest were identified: Helix 2 (residues 71-85), Helix 3 (residues 90-100), Helix 4 (residues 119-129), Helix 5 (residues 142-151), and Helix 6 (residues 156-179) (**Fig. 8B**). Of these five regions, the residues in helices 3, 4, and 5 primarily make up the negatively charged surface of IL-34 while residues in helices 2 and 6 make up IL-34 positively charged surface (**Fig. 8C**).

To further narrow down the IL-34 binding site for TREM2, we predicted TREM2/IL-34 complex structure using HADDOCK *(4)* with BLI results of TREM2 binding site for IL-34 as restraint. Our protein-protein docking results show the negatively charged surface of IL-34 (Helixes 3, 4, and 5) interacted with the positively charged basic site of TREM2 (**Fig. 8C-D**). The TREM2 binding site for IL-34 included the four key TREM2 residues of R47, R62, R76, and R77 whose mutation could greatly inhibit TREM2/IL-34 interactions when mutated as observed in BLI results. In the predicted TREM2/IL-34 complex structure (**Fig. 8D)**, TREM2 residues R47, R62, and R77 all formed hydrogen bonds with residues in IL- 34 (**Fig. 8D**). Additionally, TREM2 residue R76 formed a salt bridge with IL-34 residue D107 (**Fig. 8D**). This interaction is particularly interesting as the loss of the salt bridge when R76 is mutated to aspartic acid could be a driving factor for complete loss of binding in this mutation. Further, the mutations to R47, R62, and R77 could all reduce or inhibit the formation of key hydrogen bonds that could result in reduced interactions. In our model we also noted TREM2 residue R98 formed a salt bridge with IL-34 residue E111 (**Fig. 8D**). The mutation R98W has previously been identified in Alzheimer’s disease (AD) patients and the R98W TREM2 variant may be associated with AD [2]. The mutation of arginine to tryptophan would break the salt bridge with IL-34 and could potentially reduce TREM2/IL-34 interactions. Our computational results strongly suggest that the negatively charged surface of IL-34 (helices 3, 4, and 5) directly binds to and interacts with the TREM2 positively charged basic site.

### Competition binding experiments support that IL-34 binds to a site adjacent to those occupied by apoE4 or oAβ42

Since our binding studies indicated that IL-34 did not bind to the hydrophobic site and instead engaged a surface on the basic site centered on R76, we hypothesized that TREM2 might be able to bind to both IL-34 and either apoE4 or oAβ42 simultaneously. To determine the relationship between binding sites for IL-34, apoE4, and oAβ, we employed competition binding experiments by BLI similar to those we had done for apoE4 and oAβ42 (**Fig. 9**). First, we investigated competition between IL-34 and apoE4. We found that when TREM2 was first exposed to 500 nM apoE4 and then dipped in 500 nM IL-34, binding to IL-34 was reduced by 61.6% (**Fig. 9A,C,E**). When TREM2 was first exposed to 500 nM IL-34 and dipped into 500 nM apoE4, binding to apoE4 was reduced by 14.7% (**Fig. 9B,D,F**). These results suggest that apoE4 and IL-34 can slightly sterically inhibit one another for binding to TREM2, which likely occurs due to the proximity of R76 to the hydrophobic site. We next investigated competition between IL-34 and oAβ42. When TREM2 was first exposed to 500 nM oAβ42, binding to 500 nM IL-34 was only reduced by 18.9% (**Fig. 9G,I,K**). When TREM2 was first exposed to 500 nM IL-34, binding to oAβ42 was marginally reduced by 5.4% (**Fig. 9H,J,L**). These results suggest that IL34 and oAβ42 only slightly compete for binding to TREM2, suggesting little, if any, overlap of binding sites. Altogether, the results support that IL-34 binds a site adjacent to the hydrophobic sites engaged by apoE4 and oAβ42.

**Figure 9.**
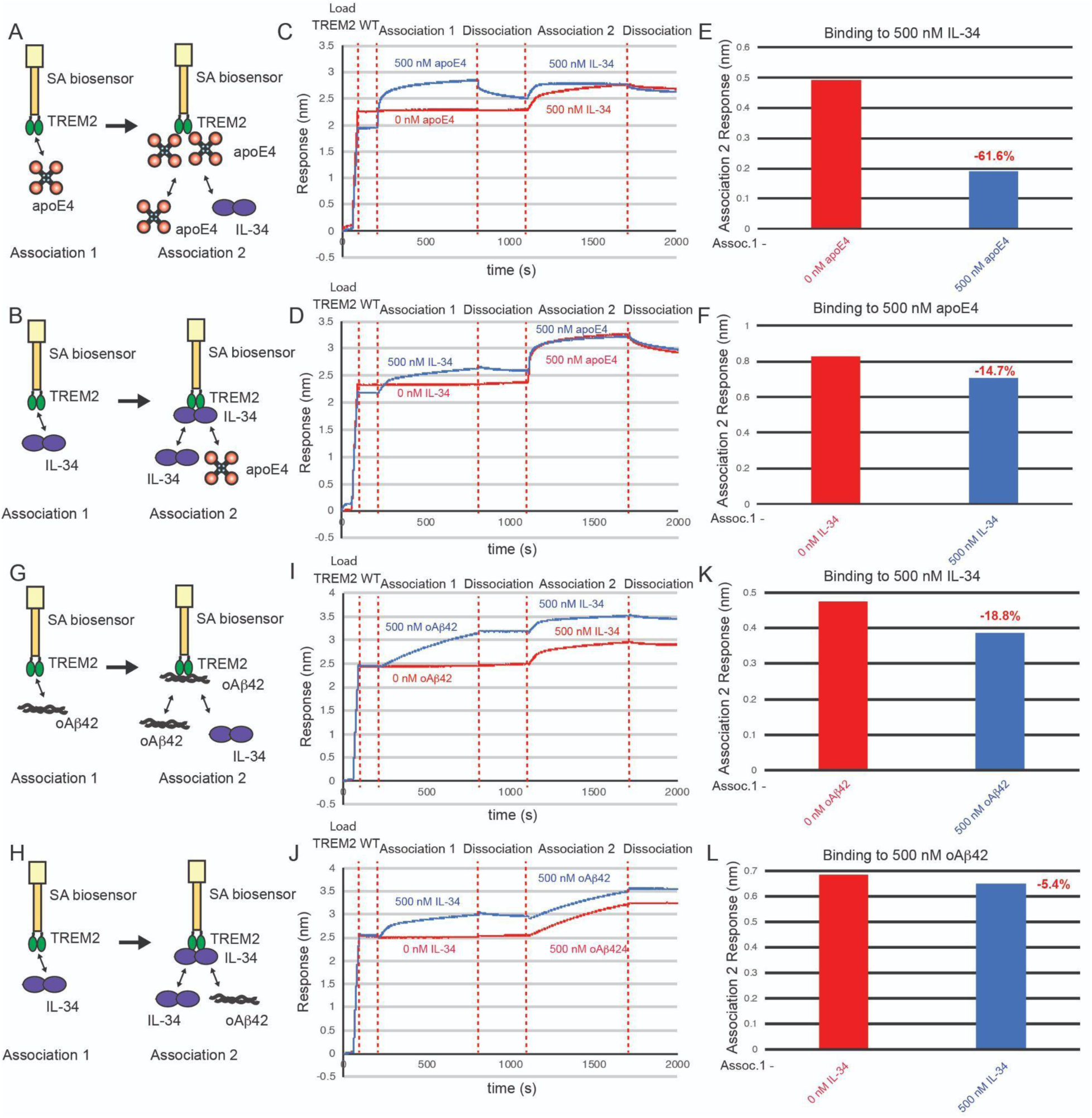
ApoE4 partially competes IL-34 for binding to TREM2, but oAβ42 does not compete IL-34 for binding to TREM2. Schematic of competition binding BLI experiments. **C&D)** BLI sensograms for **C)** apoE4 competing IL-34 binding to TREM2 and **D)** IL-34 competing apoE4 binding to TREM2. Red sensorgrams are TREM2 binding to **C)** 500 nM IL-34 or **D)** 500 nM apoE4 alone while blue sensorgrams show competition experiments where **C)** 500 nM apoE4 or **D)** 500 nM IL-34 are bound first. **E&F)** BLI binding magnitudes for TREM2 binding to **E)** 500 nM IL-34 alone or when pre-binding 500 nM apoE4 or **F)** 500 nM apoE4 alone or when pre-binding 500 nM IL-34. Percent decrease in Association 2 binding signal in the presence of the competitor is shown above the bars. **G & H)** Schematic of competition binding BLI experiments. **I&J)** BLI sensograms for **I)** IL-34 competing oAβ42 binding to TREM2 and **J)** oAb42 competing IL-34 binding to TREM2. Red sensorgrams are TREM2 binding to **I)** 500 nM IL-34 or **J)** 500 nM oAβ42 alone while blue sensorgrams show competition experiments where **I)** 500 nM oAβ42 or **J)** 500 nM IL-34 are bound first. **K&L)** BLI binding magnitudes for TREM2 binding to **K)** 500 nM IL-34 alone or when pre-binding 500 nM oAβ42 or **L)** 500 nM oAβ42 alone or when pre-binding 500 nM IL-34. Percent decrease in Association 2 binding signal in the presence of the competitor is shown above the bars.

## Discussion

TREM2 was first linked to AD by GWAS studies that identified rare point mutations in TREM2 as significant risk factors for developing AD [2, 4]. At that time, there were few known ligands for TREM2, outside of DNA and bacterial cell membrane debris [26]. Since then, a number of TREM2 ligands with relevance to AD have been identified. These ligands can all be classified as associated with tissue damage, and TREM2 should likely be classified as a scavenger receptor [34]. In this study, we begin to unravel the binding surfaces utilized by this promiscuous receptor.

The hydrophobic site on TREM2 appears to be the main binding site for most ligands. Here, we found that double and triple mutations to the hydrophobic site could ablate binding to apoE4, oAβ42, and TDP-43, whereas double mutations to the basic site or site 2 did not ablate binding to any of the studied ligands. Competition experiments between apoE4 and oAβ42 indicate that they do engage overlapping binding sites. This is in agreement with previous competitive binding experiments by BLI which also demonstrated this [15]. In that study, oAβ42 was shown to completely block apoE4 from binding to TREM2, however concentrations were not reported. In contrast, in our study we found that apoE4 could more strongly compete for binding to TREM2, with roughly a 2:1 apoE4:oAβ42 molar ratio able to completely block engagement of oAβ42 (**Fig. 5**). Conversely, oAβ42 could not strongly compete with apoE4 for binding to TREM2. The hydrophobic site is composed mostly of residues from the CDR1, CDR2, and CDR3 loops. Our molecular dynamics simulations indicate that these loops represent the most conformationally dynamic region of TREM2 [35], further suggesting this region could structurally adjust to engage diverse ligands.

The TREM2 basic site appears to be the major surface engaged by IL-34, in particular the region surrounding R76 and R77. This observation agrees with other structural studies of IL-34 in complex with its receptors. In these studies, electrostatic contacts between IL-34 and its receptors appear to drive complex formation. For example, the IL- 34 dimer contains two pronounced electronegative surfaces that pair with large electropositive surfaces on the receptor, such as CSF1R [32]. Our experiments suggest that is also the case with TREM2 binding of IL-34, and are consistent with previous demonstrations that TREM2 and CSF1R can compete for binding to IL-34 [22].

Previous studies using crosslinking as mass spectrometry suggested that site 2 was the major surface engaged by oAβ42 on TREM2 [8]. However, it should be noted in that study that the TREM2 utilized in those experiments was produced in E coli, and is therefore not glycosylated. We have noted the de- or non-glycosylated TREM2 tends to non-specifically aggregate, thus this material is not well-suited for binding studies. Here we found that double mutations at site 2 still retained binding to oAβ42, albeit in a reduced capacity. Instead, we found that double and triple mutations at the hydrophobic site could ablate binding to oAβ42. Mutations to other regions of TREM2 also lead to some decreases in binding to oAβ42, suggesting this oligomeric ligand might engage multiple sites on TREM2, with the major engagement being the hydrophobic site.

In this study, we developed a protein array of TREM2 variants that could be used to map binding to ligands. This array represents a powerful tool that can be used to map binding of TREM2 to newly discovered endogenous ligands and candidate small molecule therapeutics.

In addition to utilizing structurally designed TREM2 variants to map binding to ligands, we also investigated the major TREM2 AD risk variants R47H, R62, D87N, and T96K. In general, we have observed that the disease variants only mildly impact binding to ligands, and are not in any case loss-of-function mutations. In some instances, for example TREM2 D87N with apoE, we observe clear gain-of-function. These observations indicate that TREM2 AD variants only subtly impact interactions with ligands, and their impact is likely nuance, and will take comprehensive functional studies to understand how they are related to disease.

In summary, we find that most TREM2 ligands engage the hydrophobic site (apoE, oAβ42, TDP-43), while others bind nearby using the basic site (IL-34 and PS) (**Fig. 10**). These observations suggest that the hydrophobic and basic sites could be therapeutically targeted to modulate TREM2 function and microglia action.

**Figure 10.**
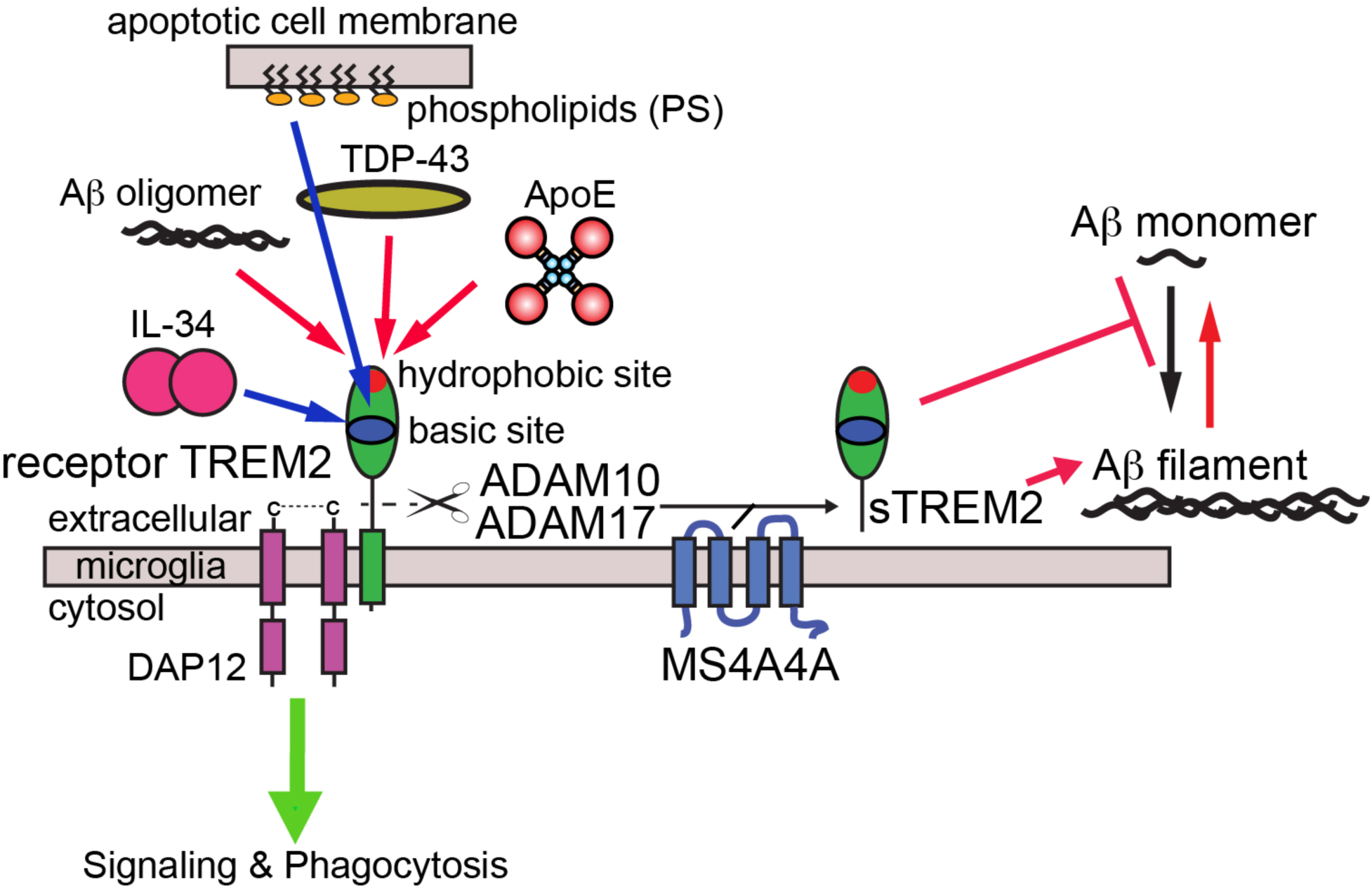
Graphic schematic for TREM2 interactions with AD ligands. Signaling interactions mapped to the basic site are denoted by blue lines, while signaling interactions mapped to the hydrophobic site are shown as red lines. The binding site for PS is from a previous crystallographic study [25], while the others are comprehensively determined in the current study. Soluble TREM2 (sTREM2) can be produced by proteolytic cleavage [36, 37] or alternative transcripts [38, 39]. Levels of proteolysis produced sTREM2 appear to be modulated by MS4A4A [40]. The sTREM2 can inhibit Aβ polymerization [8, 10] and disaggregate Aβ filaments [10].

## Acknowledgements

This work was supported by Alzheimer’s Association Research Grant (AARG-16-441560)(TJB), Bright Focus Foundation (A2022032S)(TJB), and Amgen Scholars Research Fellowships (JAG, JW).

---

**Table S1.** Binding sites on TREM2 basic site and CDR2 regions for IL-34 helices predicted by hydropathy mapping to have at least 75% percent match and a degree of complementary hydropathy (DCH) of at least 0.5.

| Region (Residues) |  | Forward |  |  |  | Reverse |  |  |  |
| --- | --- | --- | --- | --- | --- | --- | --- | --- | --- |
| IL-34 | TREM2 | Match | DCH | Start | End | Match | DCH | Start | End |
| Helix 2<br>(67-82) | Basic site<br>and CDR2<br>(62-78) | 75.00% | 0.533 | 60 | 75 | 75.00% | 0.545 | 60 | 75 |
|  | CDR2 and<br>Basic site<br>(69-78) |  |  |  |  | 75.00% | 0.54 | 71 | 86 |
|  | Basic site<br>(112-114) |  |  |  |  | 75.00% | 0.616 | 112 | 127 |
| Helix 5<br>(139-151) | Basic site<br>(47-50) | 76.92% | 0.503 | 49 | 61 |  |  |  |  |
|  | Basic site<br>(62-68) |  |  |  |  | 76.92% | 0.539 | 53 | 65 |
|  | CDR2 and<br>Basic site<br>(69-78) | 76.92% | 0.538 | 70 | 82 |  |  |  |  |
|  | Basic site<br>(112-114) |  |  |  |  | 84.62% | 0.573 | 114 | 126 |
|  | Other |  |  |  |  | 84.62% | 0.568 | 117 | 129 |
| Helix6<br>(160-181) | Basic site<br>and CDR2<br>(47-75) |  |  |  |  | 77.27% | 0.521 | 49 | 70 |
|  | Basic site<br>(112-114) | 77.27% | 0.552 | 100 | 121 |  |  |  |  |

**Table S2.**
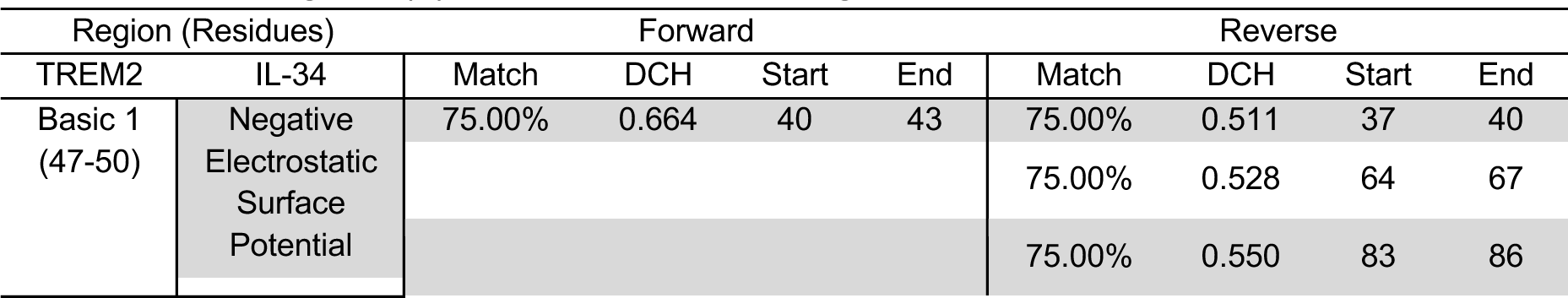
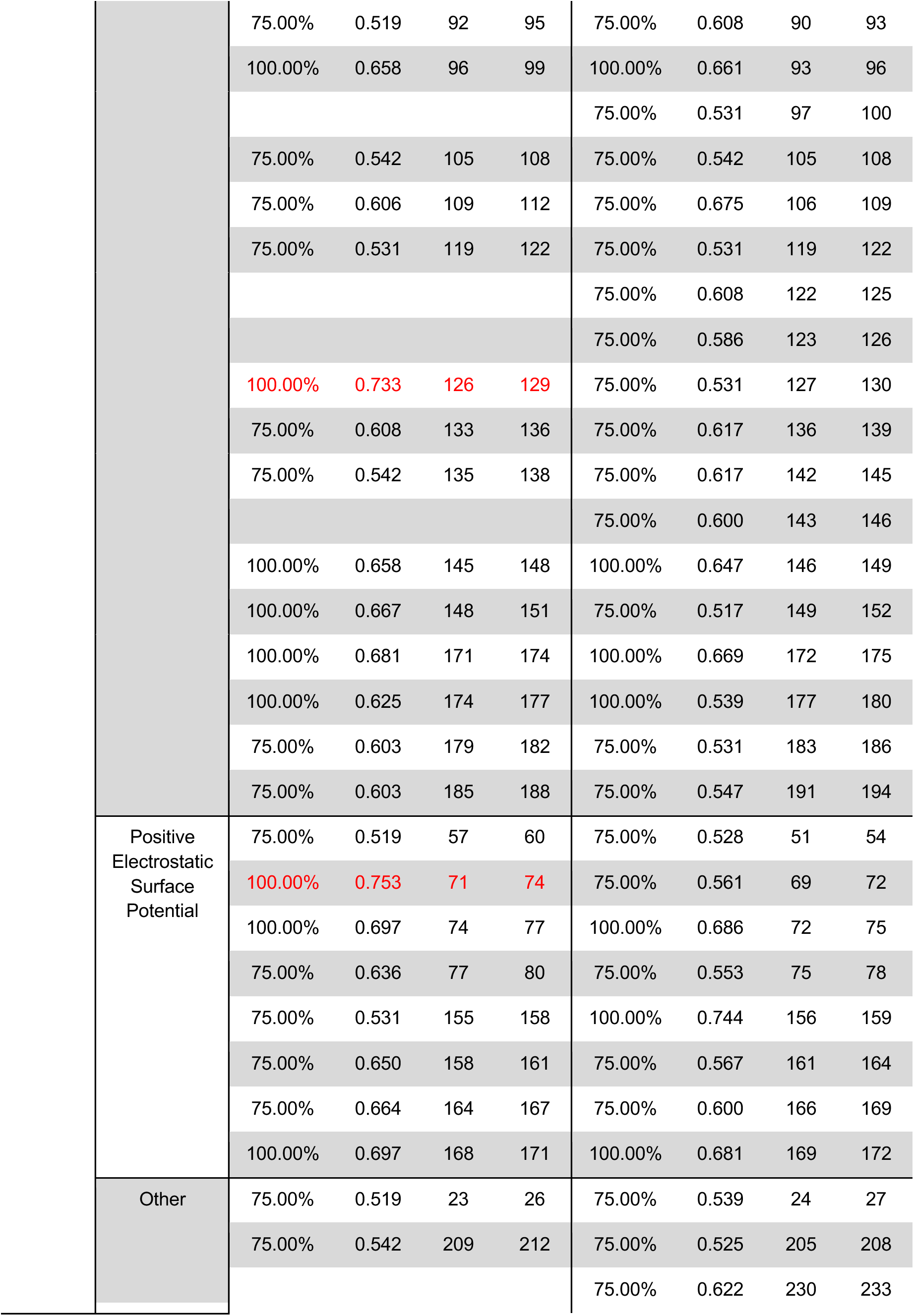
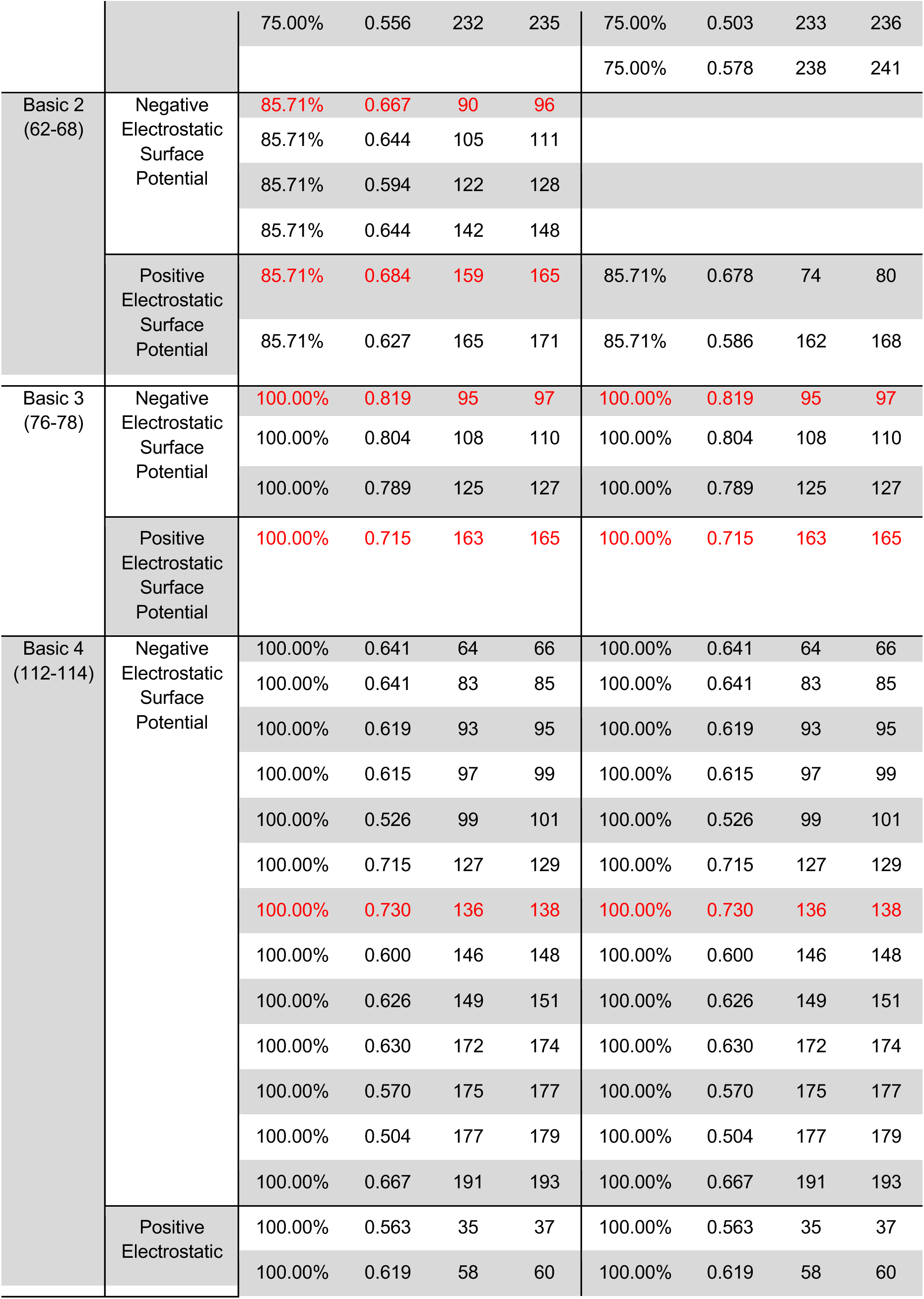
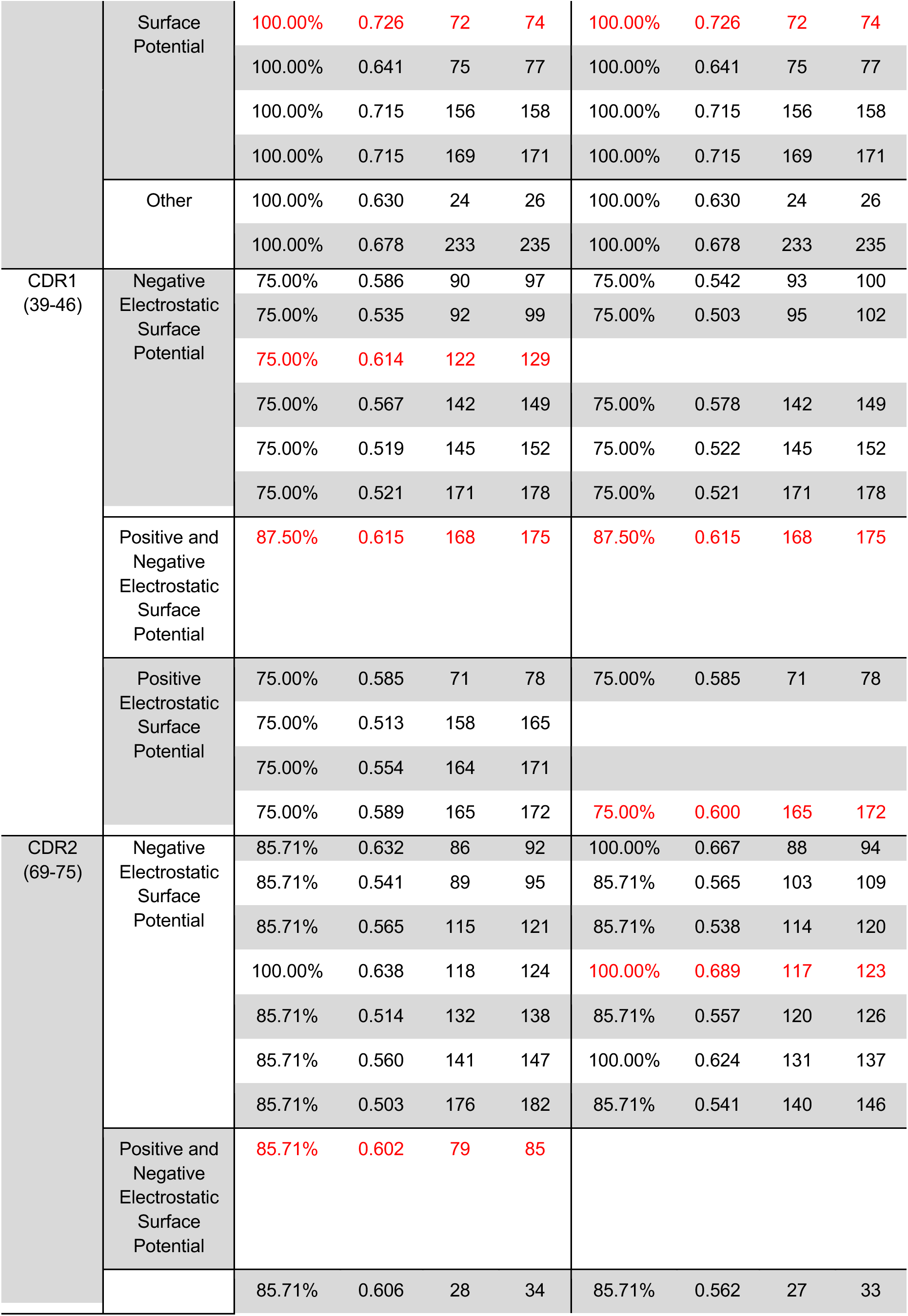
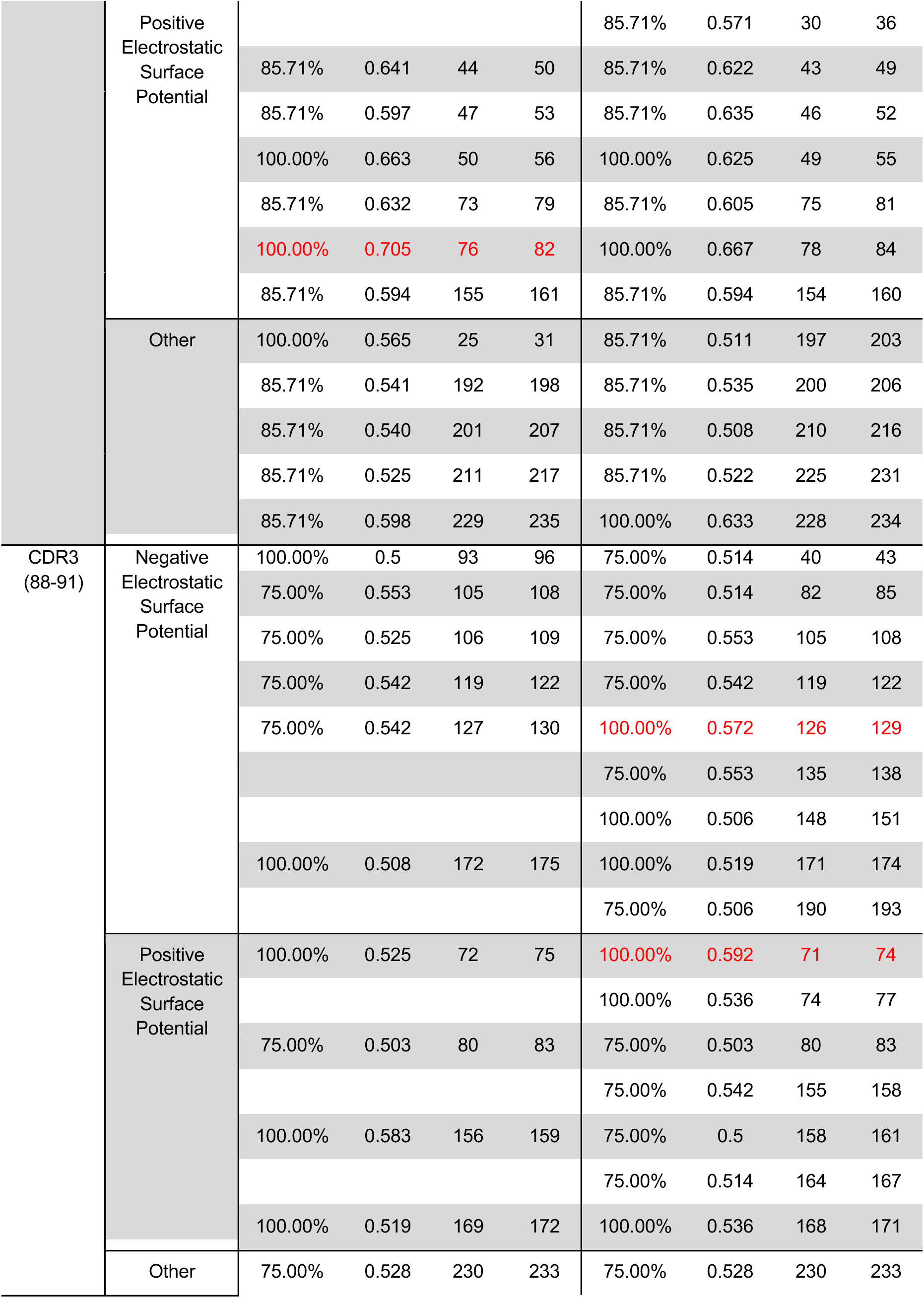
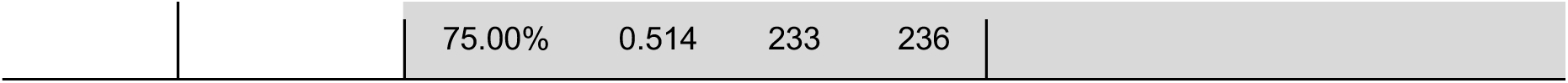
Potential binding regions on IL-34 for TREM2 predicted by hydropathy mapping to have at least 75% percent match and a degree of complementary (DCH) hydropathy of at least 0.5. Note: Rows with text colored in red represent the top predicted binding site(s) between a pair of regions.

## Notes

### Competing Interest Statement

The authors have declared no competing interest.

